# DeepDetect: deep learning of peptide detectability enhanced by peptide digestibility

**DOI:** 10.1101/2022.08.16.504211

**Authors:** Jinghan Yang, Fuzhou Gong, Yan Fu

**Affiliations:** CEMS, NCMIS, RCSDS, Academy of Mathematics and Systems Science, Chinese Academy of Sciences, Beijing 100190, China; School of Mathematical Sciences, University of Chinese Academy of Sciences, Beijing 100049, China

## Abstract

In tandem mass spectrometry-based proteomics, proteins are digested by specific protease(s) into peptides, but generally only a fraction of theoretical peptides can be detected. To explore the characteristics of proteotypic peptides, we have developed a series of methods for peptide digestibility and detectability prediction, and this is the latest one of them. Specifically, we propose here a bidirectional long short-term memory (BiLSTM)-based algorithm, named DeepDetect, for the prediction of peptide detectability enhanced by peptide digestibility. Compared with existing algorithms, DeepDetect is featured by its improved accuracy of prediction and wide applicability to commonly used proteases covering Trypsin, ArgC, Chymotrypsin, GluC, LysC, AspN, LysN, and LysargiNase. On eleven independent test data sets, DeepDetect achieved AUCs of 0.858~0.957, superior to the state-of-the-art algorithms, AP3 and PepFormer, which either used conventional machine learning methods or disregarded the digestibility feature. As an application, the predicted peptide detectability was utilized to re-rank and validate the peptide-spectrum matches, and showed the potential to promote the sensitivity of peptide identification.

## INTRODUCTION

Mass spectrometry (MS) has become the central of high-throughput proteomics, allowing several thousand proteins to be rapidly analyzed with high accuracy and low cost in a single experiment.^1,2^ In tandem MS-based proteomics, proteins are digested by specific protease(s) into peptides to generate tandem mass spectra with intricate information regarding amino acid sequences. However, only a fraction of theoretical peptides can be generally detected (or observed) during the experiments, inhibiting comprehensive characterization of proteins in the sample. Thus, peptide detectability, defined as the probability of a peptide being observed from a standard sample in a standard proteomics routine, has become a crucial feature for proteome analysis.^3^ Unfortunately, the mechanism of peptide detection is complicated on account of several factors, such as protein digestion efficiency, ionization efficiency, post-translational modifications, etc.^4^

In targeted quantitative proteomics, proteotypic peptides (PTPs), i.e., detectable and unique representatives of proteins, are commonly measured to quantify target proteins of interest. Although the selection of PTPs can be managed by experimental information along with expert rules, the detectability of numerous peptides still cannot be assessed due to incomplete experimental data.^4^ Therefore, researchers have turned to computational approach for accurate prediction of peptide detectability. In addition, the Human Proteome Project (HPP)^5,6^ teams have been dedicated to detecting the proteins without sufficient MS evidence of their existence, also referred to as “missing proteins”. Recently, it was reported that the predicted detectability of unique tryptic peptides from the missing proteins were significantly low.^7^ This result indicated that the peptide detectability problem could be a reason for the difficult detection of missing proteins. Consequently, the peptide detectability feature could be an indicator to assess the quality of identified results by one search engine.

Till now, various studies have been conducted to characterize the mechanism of peptide detection to achieve accurate prediction of peptide detectability. For example, Guruceaga et al.^7^ evaluated the performance of different machine learning (ML) methods with over 550 features, mainly originating in amino acid sequences and properties from the AAindex database. It turned out that the random forest classifier with nonredundant features exhibited higher accuracy than other methods. Besides, Gao et al.^8^ presented an Advanced Proteotypic Peptide Predictor (AP3), which initiatively incorporated the predicted peptide digestibility as a significant feature for the prediction of proteotypic peptides, considering that protein digestion is a key step before MS analysis. In contrast with conventional ML methods, deep learning (DL) is more adept at feature extraction and tends to learn more efficacious representations from raw data, such as sequence, image, etc. Hence, Zimmer et al.^4^ introduced a deep neural network based method, d::pPop, to reformulate the PTP prediction problem into a rank regression problem instead of a binary classification problem with the converted features. Specifically, d::pPop managed to rank the peptides within an interested protein to select PTPs with the maximal detectability for the design of targeted proteomics quantification assays. Afterwards, Serrano et al.^9^ implemented a DL tool based on convolutional neural network using amino acid sequences alone, called DeepMSPeptide, to acquire better performance of peptide detectability prediction than previous studies^4,7,10^.

Lately, Cheng et al.^11^ built PepFormer, an end-to-end Siamese network based on the Transformer and gated recurrent unit, to predict peptide detectability by amino acid sequences only. Subsequently, Yu et al.^12^ applied capsule network with convolutional block attention module to predict the detectability of peptides by extracting the peptide chain features, covering the residue conical coordinate, the amino acid composition, the dipeptide composition, and the sequence embedding code.

Although several DL tools have been provided with better performance than ML, they still have some defects. First, it is known that the major characteristic of DL as a kind of end-to-end algorithm, is the ability to directly extract information from raw data. But some of the DL methods still depended on hand-crafted features, leading to insufficient learning.^4,12^ Second, the protein digestion, as a key step in the proteomic workflow, is never complete (existence of missing cleavages). However, the existing DL methods did not consider the significant influence of digestion efficiency and heterogeneity on peptide detection. Third, there is still a lack of versatile tools to predict peptide detectability for a variety of proteases.

Here, we propose an extended deep learning algorithm, named DeepDetect, to accurately predict peptide detectability for eight commonly used proteases, covering Trypsin,^13–17^ ArgC,^13,17^ Chymotrypsin,^13,17^ GluC,^13,17^ LysC,^13,17^ AspN,^13,17^ LysN,^13,18^ and LysargiNase.^19,20^ First, the bidirectional long short-term memory (BiLSTM) network is used to extract the context features among amino acids. Next, the probability predicted by BiLSTM is calibrated with the peptide digestibility predicted by DeepDigest.^21^ The area under the ROC curves (AUCs) of DeepDetect on eleven independent test data sets were 0.858~0.957, superior to the state-of-the-art ML and DL tools, i.e., AP3^8^ and PepFormer.^11^ Furthermore, the accuracies of DeepDetect were significantly improved by peptide digestibility, compared to DeepDetect without this feature. As an application of DeepDetect, the predicted peptide detectability was served as a supplementary feature in Percolator^22^ to re-rank and validate the peptide-spectrum matches (PSMs). The results showed that peptide detectability was a helpful feature to promote the sensitivity of peptide identification.

## EXPERIMENTAL PROCEDURES

### Data sets

DeepDetect was first trained on eight large-scale training data sets, involving eight commonly used proteases, i.e., Trypsin, ArgC, Chymotrypsin, GluC, LysC, AspN, LysN, and LysargiNase (Table 1).^13,19^ Next, the model was carefully evaluated on eleven test data sets from samples of E.coli, yeast, mouse, and human.^14–18,20^ All of these data sets except 2012Heck_LysN_Yeast were also utilized in DeepDigest.^21^ Specifically, we chose a bigger data set (2012Heck_LysN_Yeast) for LysN protease in this study.

**Table 1.**
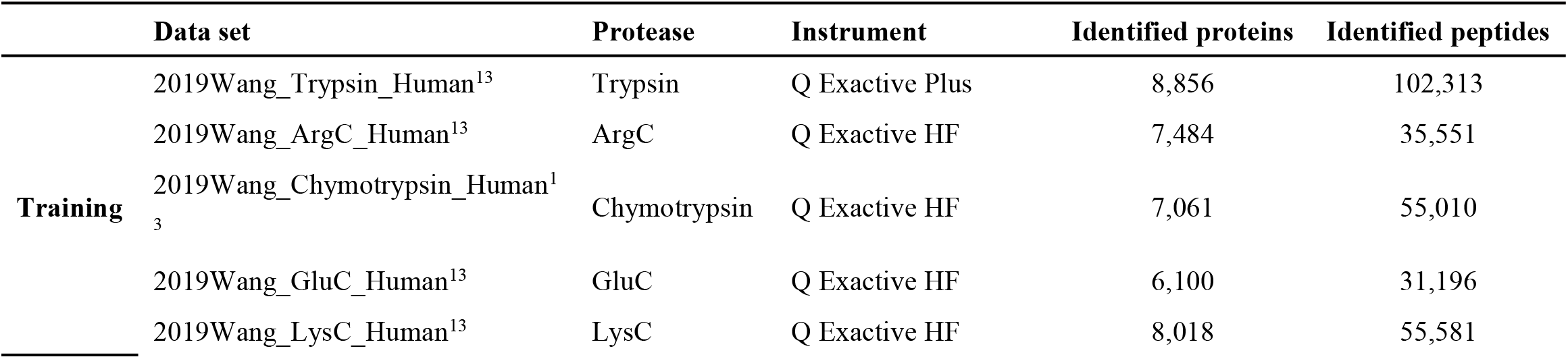
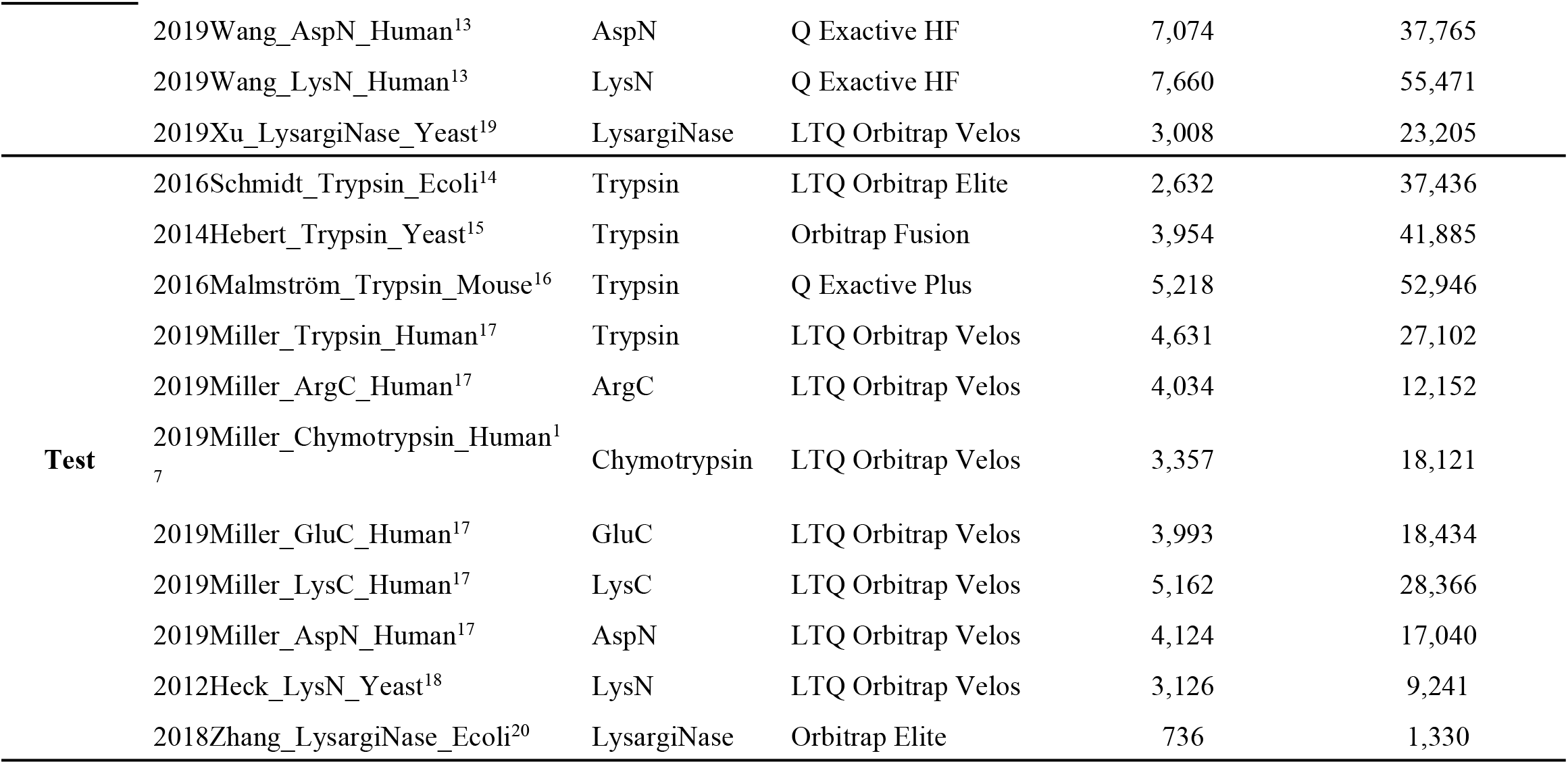
Public data sets used for training and test.

The raw files were publicly available and downloaded from PRIDE. They were searched by MaxQuant^23,24^ with the same parameters as in our previous study.^21^ Except for the yeast data set from Heck’s lab,^18^ precursor mass tolerance was set to 50 ppm for the first search and 4.5 ppm for the main search; fragment mass tolerance was set to 0.5 Da; the variable modifications were acetylation on protein N-term, oxidation on methionine, and phosphorylation on serine and threonine; the fixed modification was carbamidomethylation on cysteine; and the false discovery rate (FDR) was set to 1% at both peptide and protein levels.

### Data Preprocessing

At first, we sorted the identified proteins by spectral counts (SCs) and sequence coverages respectively in descending order. Intuitively, the highly ranked proteins tend to be more confident. Thus, identified proteins in the top 50% of both ranks were then collected to construct our data sets.^8^ For each protein, in silico digestion was carried out with up to two missed cleavages and the same length range as the identified peptides. At last, peptides with *SC >* 1 were labeled as positive samples, while the unidentified peptides were labeled as negative samples.

### Workflow of DeepDetect

As illustrated in Figure 1, DeepDetect is mainly based on the BiLSTM network, and further enhanced by peptide digestibility.^21^ Peptide sequence with variable length is the only input. After zero padding, the embedding layer is used to encode the sequence into a matrix of shape *l_max_* × 10. *l_max_* is the maximum length allowed for all the input peptides, which can be self-defined. To capture full information of peptide sequence, *l_max_* is set to the maximum length of all the input peptides. Subsequently, BiLSTM is used to extract a 40-dimensional feature vector. Finally, the probability (*p_BiLSTM_*) is predicted by a sigmoid function as the output.

**Figure 1.**
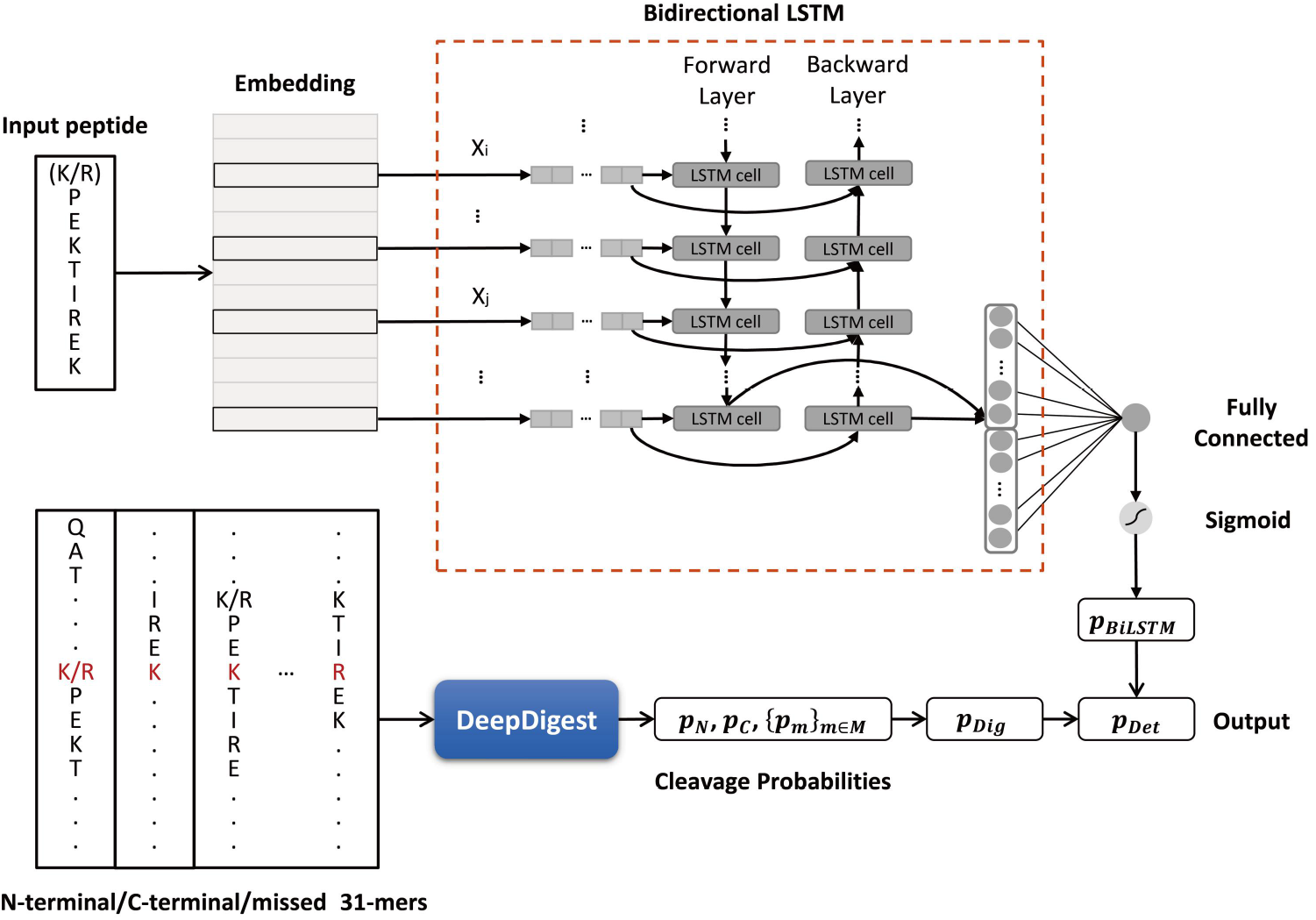
Workflow of DeepDetect.

Meanwhile, the cleavage probabilities of potential cleavage sites are predicted by DeepDigest.^21^ Then, the peptide digestibility is calculated by the following formula:

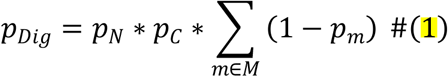

where *p_N_* and *p_c_* are the predicted cleavage probabilities of the N- and C-terminal sites of peptide, respectively; *p_m_* is the predicted cleavage probability of each missed cleavage site in the peptide, and *M* is the set of all the missed cleavage sites in the peptide.

Eventually, peptide detectability is calculated as follows:

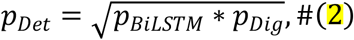

where *p_BiLSTM_* is the predicted probability of BiLSTM network, and *p_Dig_* is the predicted peptide digestibility of DeepDigest.

DeepDetect was implemented in Python (v3.5.2), using Keras (v2.2.4, https://keras.io/) with TensorFlow^25^ (v1.10.0) backend. The models were trained for 50 epochs with batch size of 100 by ADAM optimizer. Dropout technique was used to avoid overfitting, whose rate was selected by 3-fold cross-validation.^26^ Besides, down-sampling technique was adopted to obtain balanced data sets, in which the number of positive and negative samples were the same.

## RESULTS AND DISCUSSIONS

### Performance Evaluation and Enhancement by Peptide Digestibility

Four metrics were utilized to evaluate the performance of DeepDetect, including AUC, accuracy (ACC), Matthews correlation coefficient (MCC), and F1 score. The model was first trained on eight training data sets for each protease, and then tested on eleven independent test data sets (Table 1). The test AUCs were 0.858~0.957 as shown in Figure 2, while the test ACCs, MCCs and F1 scores were 0.773~0.889, 0.547~0.780 and 0.775~0.893, respectively, as shown in Table 2. These results demonstrated that DeepDetect had the advanced generalization capability for a variety of proteases.

**Figure 2.**
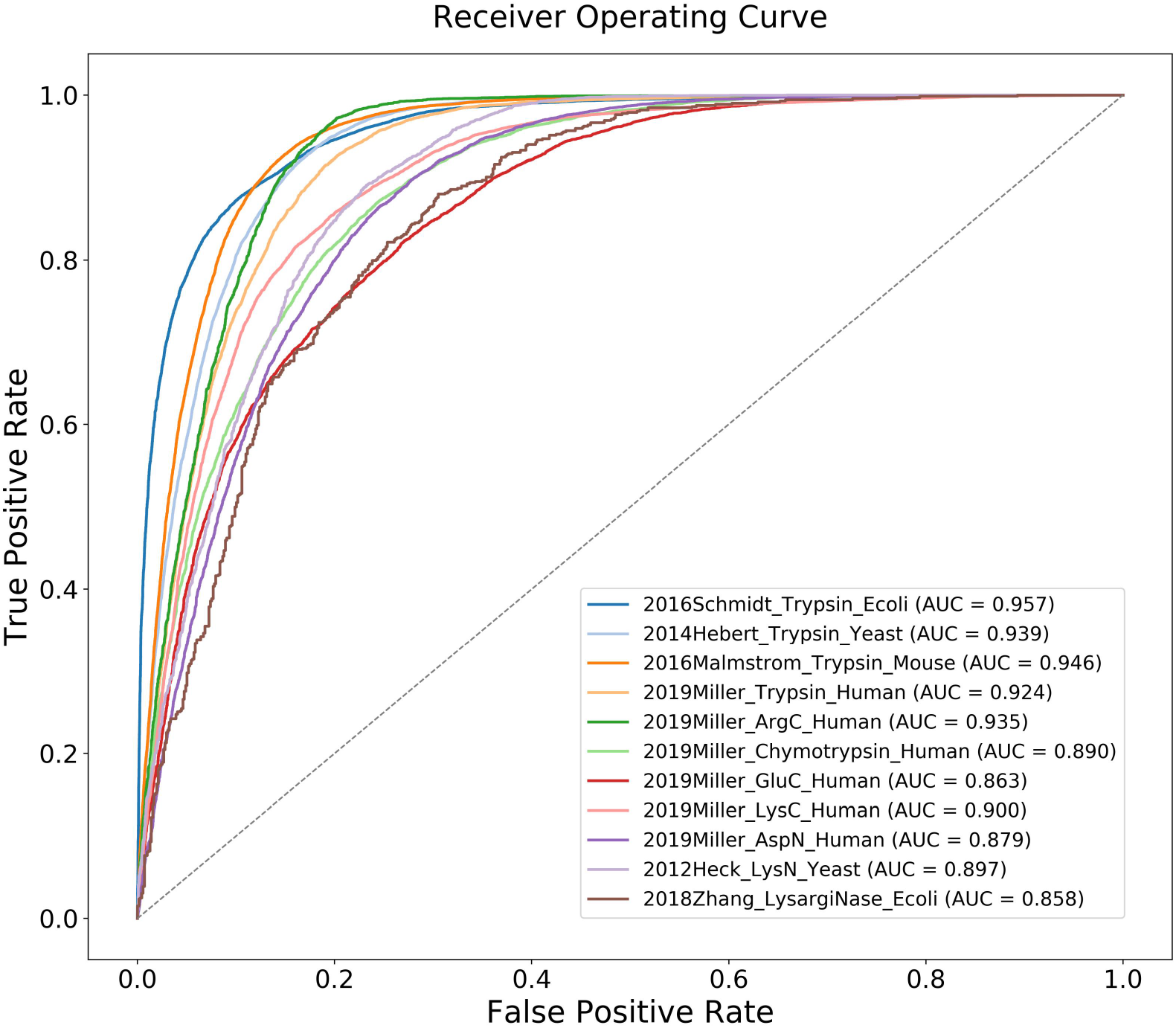
Test AUCs of DeepDetect on eleven independent test data sets.

**Table 2.**
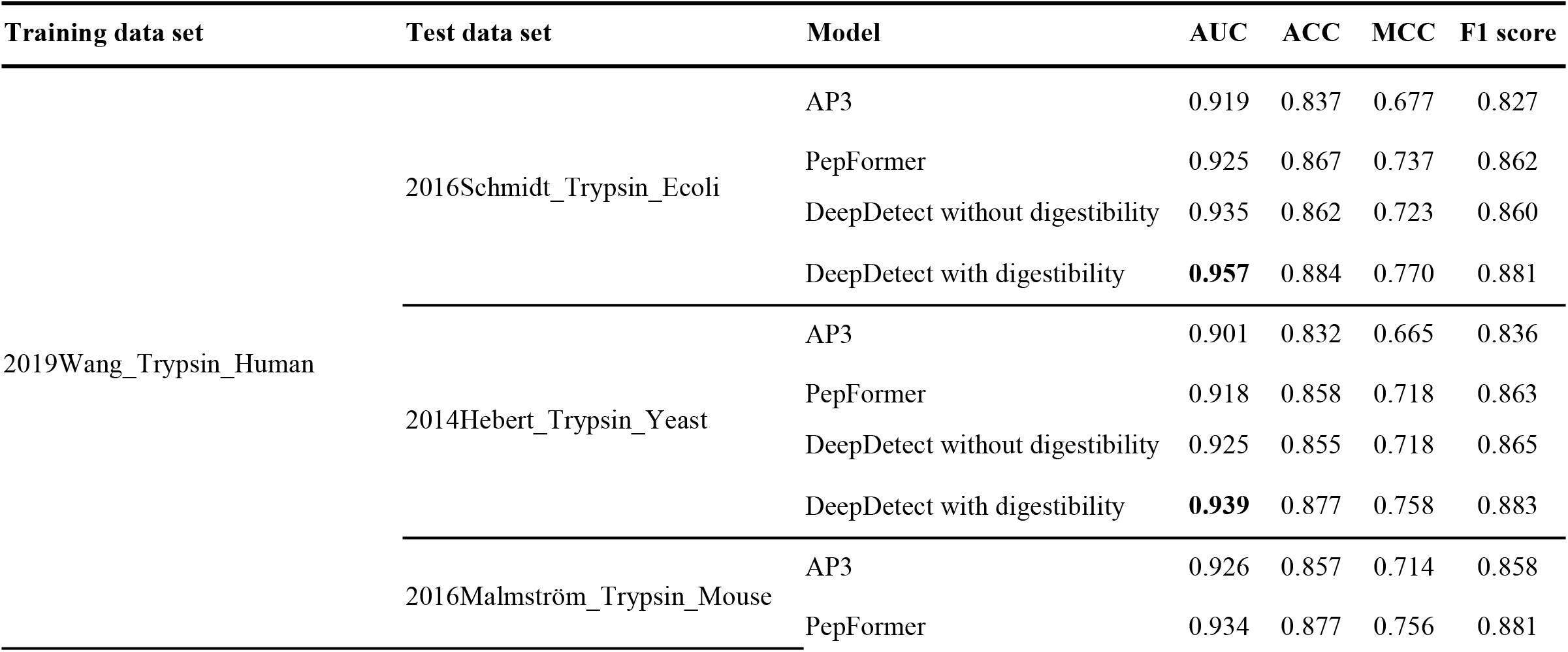
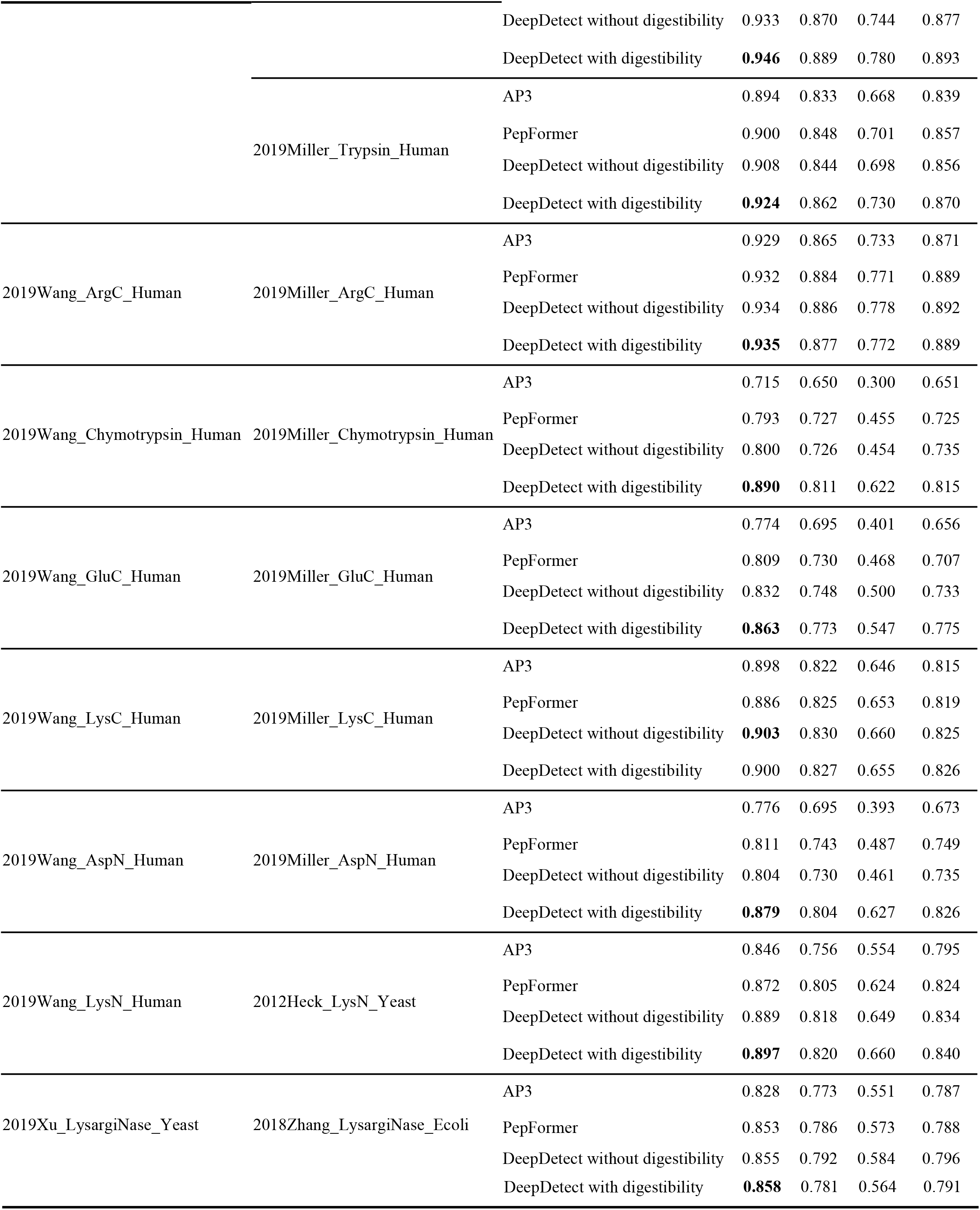
Performance comparison of AP3, PepFormer and DeepDetect (without and with peptide digestibility) on eleven independent test data sets.

Generally, proteins are first digested by specific protease(s) into peptides and then analyzed by liquid chromatography-tandem MS in a shotgun proteomics routine. Consequently, the pattern of protein digestion has a significant influence on peptide detection, with the knowledge of which peptides could be digested out and subjected to subsequent analysis. In our previous study,^8^ peptide digestibility has been proven to be an important feature for accurate prediction of peptide detectability. Naturally, this information should be taken into consideration.

To explore the impact of peptide digestibility, the performances of DeepDetect with and without this feature were compared on eleven independent test data sets. As shown in Table 2, the AUCs of DeepDetect were effectively improved by peptide digestibility. Particularly, on the 2019Miller_Chymotrypsin_Human test data set, the AUC, ACC, MCC, and F1 score increased by 11.3%, 11.7%, 37.0%, and 10.9%, respectively, with this feature considered. These results confirmed that peptide digestibility substantially enhanced the performance of peptide detectability prediction.

### Comparison with Other Tools

To further evaluate the performance of our algorithm, DeepDetect was compared with the state-of-the-art ML and DL tools for peptide detectability prediction, i.e., AP3^8^ and PepFormer^11^. Since both methods were not proposed for other proteases than trypsin, we first extended AP3 for eight proteases and retrained PepFormer on the same eight training data sets as DeepDetect, and then assessed them on eleven independent test data sets (Table 1). As a conventional ML method, AP3 exhibited competitive performance compared to PepFormer as shown in Figure 3. The AUCs, ACCs, MCCs and F1 scores were shown in Table 2. Overall, the AUCs of DeepDetect were higher than both AP3 (0.715~0.926) and PerFomer (0.793~0.934), the ACCs, MCCs and F1 scores were superior or comparable to them. These results suggested the superiority and versatility of DeepDetect.

**Figure 3.**
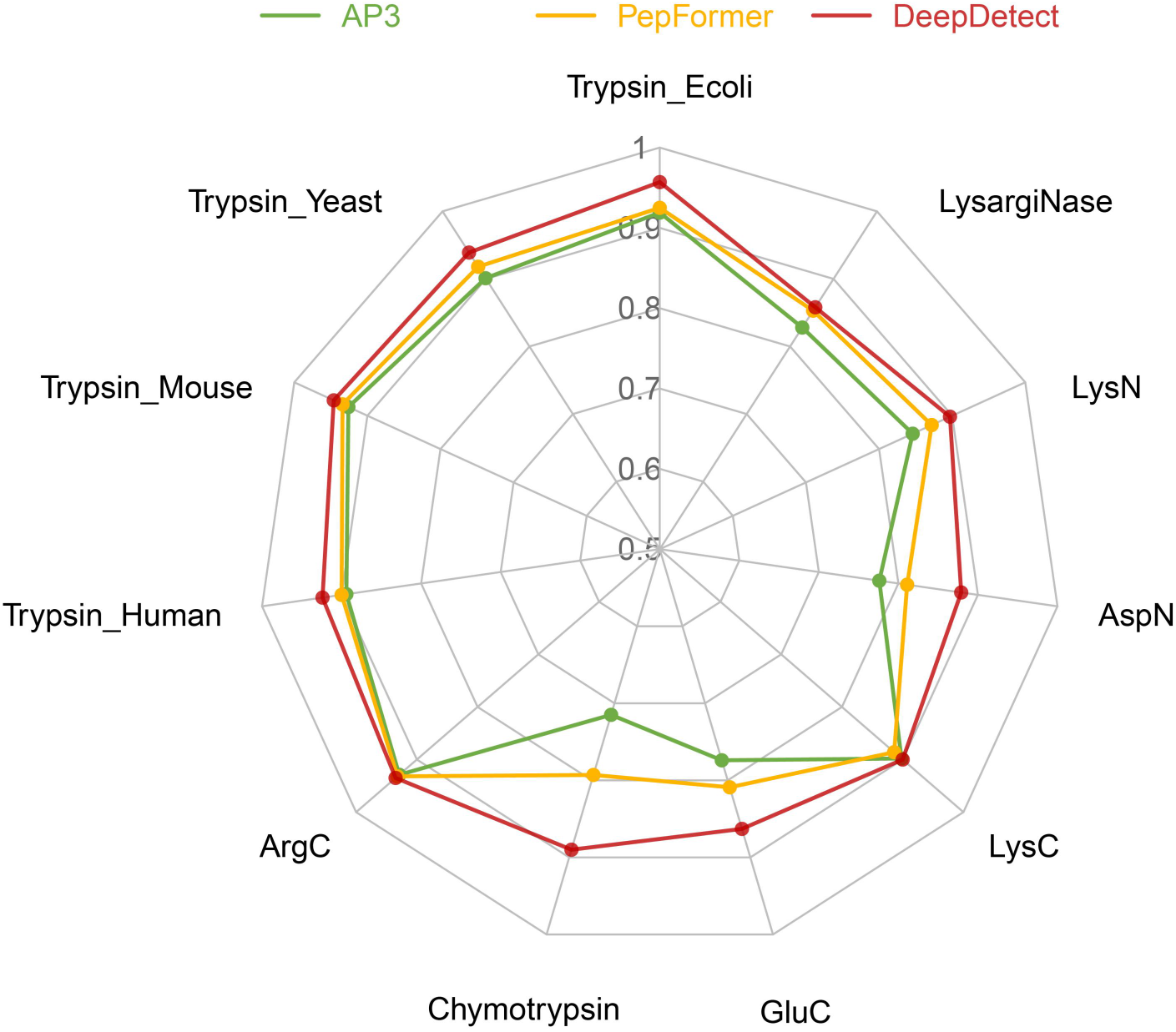
Performance comparison of AP3, PepFormer and DeepDetect on eleven independent test data sets.

### An Application of DeepDetect

In our previous study,^21^ peptide digestibility efficaciously improved the sensitivity of peptide identification. Considering that peptide detection is another crucial process in a typical shotgun proteomics experiment after protein proteolytic digestion,^8^ it is sensible to speculate that peptide detectability is also a useful feature for PSM validation. To substantiate this speculation, Percolator (version 3.05.0) was utilized to re-rank the PSMs of Comet with various feature combinations.

The raw files of the four tryptic test data sets were first searched by Comet search engine^27^ in Crux^28^ pipeline with the same parameters in their original papers.^14–17^ For each (target or decoy) peptide of the top PSMs, the predicted detectability was calculated following formula (1-2). Afterwards, two pin files with and without the peptide detectability feature were generated and re-ranked by Percolator. As shown in Table 3, in contrast with the results re-ranked by the original features of Percolator (Original), the numbers of peptide identifications with peptide detectability (*p_Det_*) were boosted by 0.26%, 0.39%, 1.07%, and 0.81%, respectively, on the four tryptic test data sets under 1% FDR. These results showed that the peptide detectability predicted by DeepDetect indeed had the potential of promoting the sensitivity of peptide identification.

**Table 3.**
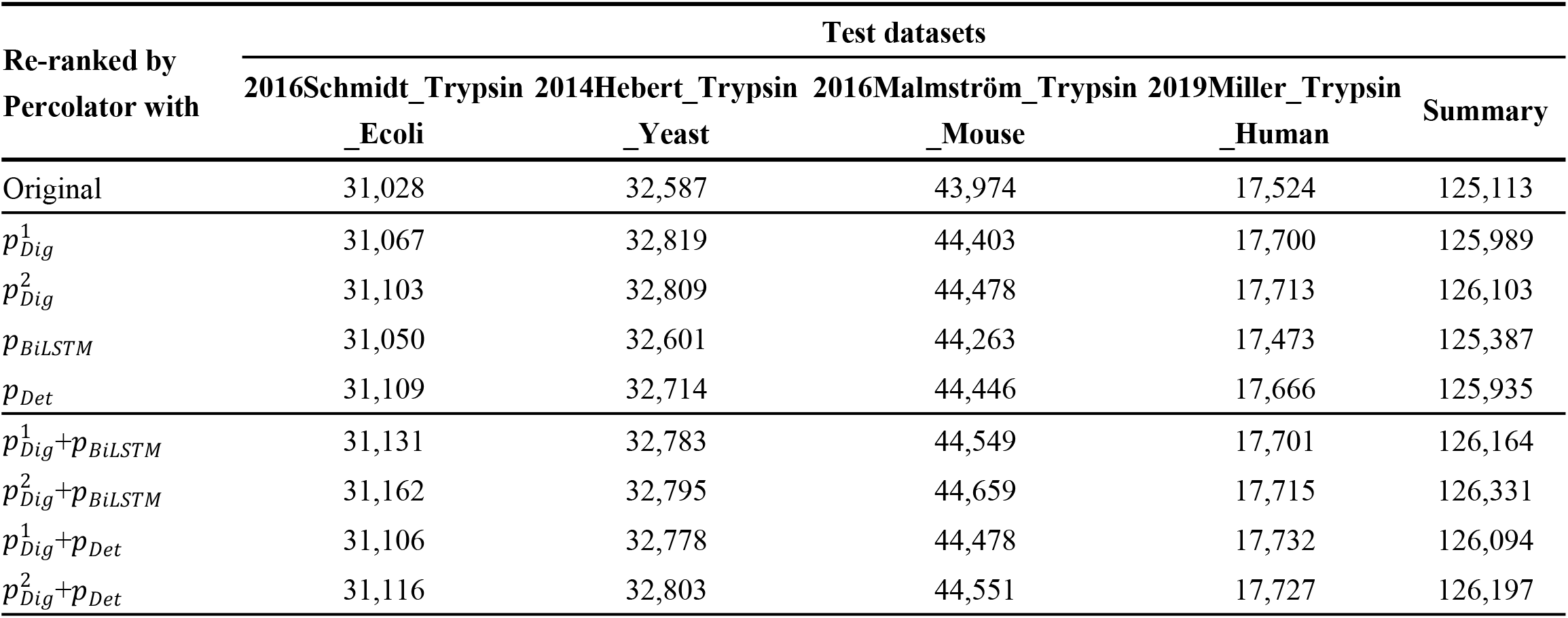
Identifications under 1% FDR (at the peptide level) after Percolator re-ranking with various feature combinations.

| Re-ranked by<br>Percolator with | Test datasets |  |  |  | Summary |
| --- | --- | --- | --- | --- | --- |
|  | 2016Schmidt_Trypsin<br>_Ecoli | 2014Hebert_Trypsin<br>_Yeast | 2016Malmström_Trypsin<br>_Mouse | 2019Miller_Trypsin<br>_Human |  |
| Original | 31,028 | 32,587 | 43,974 | 17,524 | 125,113 |
| $p_{Dig}^1$ | 31,067 | 32,819 | 44,403 | 17,700 | 125,989 |
| $p_{Dig}^2$ | 31,103 | 32,809 | 44,478 | 17,713 | 126,103 |
| $p_{BiLSTM}$ | 31,050 | 32,601 | 44,263 | 17,473 | 125,387 |
| $p_{Det}$ | 31,109 | 32,714 | 44,446 | 17,666 | 125,935 |
| $p_{Dig}^1 + p_{BiLSTM}$ | 31,131 | 32,783 | 44,549 | 17,701 | 126,164 |
| $p_{Dig}^2 + p_{BiLSTM}$ | 31,162 | 32,795 | 44,659 | 17,715 | 126,331 |
| $p_{Dig}^1 + p_{Det}$ | 31,106 | 32,778 | 44,478 | 17,732 | 126,094 |
| $p_{Dig}^2 + p_{Det}$ | 31,116 | 32,803 | 44,551 | 17,727 | 126,197 |

Moreover, to thoroughly examine the significance of these features in PSM validation, eight feature combinations of peptide digestibility and detectability were further compared, including 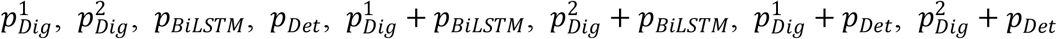. Specifically, two primary formulas were applied to calculate peptide digestibility as follows,

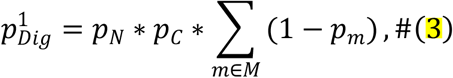

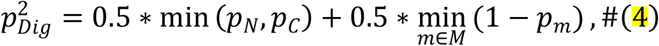

More unique peptides were identified with 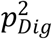 than with 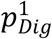 as shown in Table 3, which was the reason we chose 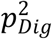 in our previous study.^21^ Besides, *p_Det_* was enhanced by peptide digestibility and naturally had a bigger influence than *p_BiLSTM_*, ensuring more identifications. However, the numbers of identified peptides with 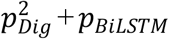 were the highest, rather than with 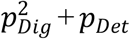. In contrast with the original results, the numbers of peptide identifications with 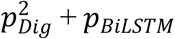 were boosted by 0.43%, 0.64%, 1.55%, and 1.09%, respectively. With the consideration of the peptide digestibility feature in 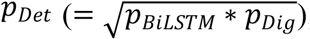, the feature combination of 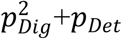 contained redundant information (*p_Dig_*), which could be the reason for the inferior performance to 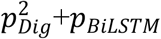. In conclusion, *p_Det_* enhanced by 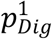 was a promising feature for PSM validation, and 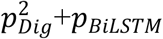 was the preferable feature combination to raise the number of identified peptides.

### Software and Data Availability

The DeepDetect tool and test data can be downloaded at http://fugroup.amss.ac.cn/software/DeepDetect/DeepDetect.html.

## CONCLUSIONS

In this study, we proposed a deep learning-based algorithm, named DeepDetect, to predict peptide detectability for eight commonly used proteases. The independent test results demonstrated the advanced generalization capability of DeepDetect. The improved performance of DeepDetect is largely owed to the inclusion of the peptide digestibility feature predicted by the DeepDigest tool we previously developed. In comparison with AP3 and PepFormer, DeepDetect exhibited higher accuracies than these state-of-the-art tools, which either used conventional machine learning methods or disregarded the digestibility feature. In the end, the application of DeepDetect showed that the predicted peptide detectability was a promising feature for PSM validation. In addition to targeted proteomics and missing protein detection, this feature could also be beneficial to biomarker discovering in cancer research^29^ and data-independent acquisition proteomics.

## ACKNOWLEDGEMENTS

This work was supported by the National Natural Science Foundation of China [No. 32070668].

